# A new LD protein, ApoL6 disrupts the Perilipin 1-HSL interaction to inhibit lipolysis

**DOI:** 10.1101/2022.11.10.516022

**Authors:** Yuhui Wang, Hai P Nguyen, Pengya Xue, Ying Xie, Danielle Yi, Frances Lin, Jose A Viscarra, Nnejiuwa U Ibe, Robin E Duncan, Hei Sook Sul

## Abstract

ApoL6 is a new LD-associated protein containing an apoprotein-like domain, expressed mainly in adipose tissue, specifically in adipocytes. ApoL6 expression is low in fasting but induced upon feeding. ApoL6 knockdown results in smaller LD with lower triglyceride (TAG) content in adipocytes, while ApoL6 overexpression causes larger LD with higher TAG content. We show that ApoL6 effect in adipocytes is by inhibition of lipolysis. While ApoL6, Perilipin 1 (Plin1) and HSL can form a complex on LD, C-terminal domain of ApoL6 directly interacts with Plin1, to compete with Plin1 binding to HSL through Plin1 N-terminal domain, thereby keeping HSL in a “stand by” status. Thus, ApoL6 ablation decreases WAT mass, protecting mice from diet-induced obesity, while adipose overexpression increases WAT mass to bring obesity and insulin resistance with hepatosteatosis, making ApoL6 a potential future target against obesity and diabetes.

## INTRODUCTION

White adipose tissue (WAT) plays a central role in energy metabolism by storing excess energy as triglycerides (TAG). Adipocytes contain a large unilocular lipid droplet (LD) and are composed of core neutral lipids, TAG and some cholesterol ester, surrounded by a single phospholipid monolayer. During periods of energy deprivation, hydrolysis of TAG stored in WAT is stimulated to release fatty acids (FA) into circulation so that other organs can use them as energy source. Therefore, lipolysis within adipocytes represents a critical process in WAT’s function of FA release and needs to be regulated exquisitely according to nutritional conditions. In modern society, however, obesity characterized by increased adipose tissue mass has become an epidemic and is associated with chronic metabolic diseases, such as type 2 diabetes and insulin resistance. Furthermore, obesity-associated ectopic TAG storage in other tissues, such as liver, contributes to insulin resistance (Unger and Scherer 2010).

Lipolysis proceeds in an orderly and regulated manner, by converting triacylglycerol (TAG) to diacylglycerol (DAG), monoacylglycerol (MAG) and FAs. These reactions catalyzed by adipose triglyceride lipase (ATGL, PNPLA2, Desnutrin) (Zimmermann et al. 2004; Villena et al. 2004; Jenkins et al. 2004; Ahmadian et al. 2009; Duncan et al. 2010), hormone-sensitive lipase (HSL) (Haemmerle et al. 2002), and monoglyceride lipase (MGL) (Taschler et al. 2011), respectively. Recent advancement in understanding lipolysis has revealed various dynamic regulatory processes: Posttranslational modifications of lipases and LD-associated proteins by hormonal signals, such as catecholamines and insulin, have opposing effects on lipolysis. Phosphorylation of lipases and LD-associated proteins regulate assembly and disassembly of protein complexes for lipolysis on the surface of LD, thereby controlling lipases to access TAG and their catalytic activities (Granneman et al. 2007; Granneman et al. 2009; Sztalryd et al. 2003; Yamaguchi et al. 2004). In the fasted or stimulated state, hormones initiate signaling cascades that increase PKA activity to activate lipolytic pathways in adipocytes. For example, HSL is phosphorylated at S563 and S660 by PKA. ATGL is phosphorylated at S406 by AMPK which has been shown to be activated by cAMP pathway (Ahmadian et al. 2011; Yin, Mu, and Birnbaum 2003; Kim et al. 2016). In addition, PKA phosphorylates many of the components of the lipolytic complex. Perilipin 1 (Plin1, Perilipin A), a LD-associated protein, having six Protein Kinase A (PKA) consensus phosphorylation sites (R(R/K)XS), is phosphorylated by PKA (Garcia et al. 2003). In addition, a cofactor of ATGL, α/β hydrolase domain–containing protein 5, CGI-58 (ABHD5), is phosphorylated at S239 by PKA (Sahu-Osen et al. 2015). Phosphorylation initiates the translocation of lipases from the cytosol to LD, enabling protein-protein interaction to assemble the lipolytic complex on the Plin1 scaffold on LD (Miyoshi et al. 2007; Miyoshi et al. 2006). Specifically, phosphorylation of Plin1 releases CGI-58 upon its phosphorylation. Then, the Phosphorylated CGI-58 (P-CGI-58) interacts with phosphorylated ATGL for TAG hydrolysis (Granneman et al. 2007; Granneman et al. 2009; Shin et al. 2017). Thus, ATGL hydrolytic activity increases by more than 20-fold upon interaction of ATGL with P-CGI-58. Phosphorylated HSL translocates from the cytosol to the surface of LD, where it binds to P-Plin1 to hydrolyze primarily DAG (Clifford et al. 2000; Egan et al. 1992). In fact, HSL translocation requires phosphorylation of both HSL and Plin1 (Haemmerle et al. 2002; Osuga et al. 2000; Su et al. 2003). Finally, MGL cleaves the remaining fatty acid (FA) from glycerol backbone (Karlsson et al. 1997). In the fed state, insulin released from pancreatic islet β cells activates phosphodiesterase 3B to decrease cAMP levels to decrease PKA activity and insulin also activates protein phosphatase 1 (PP-1) (Begum 1995; Brady and Saltiel 2001) to potentially dephosphorylate target proteins, such as Plin1 and HSL, resulting in suppression of lipolysis in adipocytes (Begum 1995; Clifford et al. 1998) Human and rodent genetic studies corroborate this concept of lipid mobilization pathway in adipocytes and the components of lipolytic pathway may serve as therapeutic targets (Schweiger et al. 2009; Schweiger et al. 2017; Yang and Mottillo 2020). In addition, there are other LD-associated proteins that may regulate this lipolytic process. In this regard, G0/G1 switch gene 2 (G0S2) has been identified as an inhibitor of ATGL by binding with the catalytic patatin domain and suppressing activation (Yang et al. 2010).

Here, in an attempt to better understand the role of LD-associated proteins on TAG metabolism of WAT, we identify an adipose-specific LD-associated protein, ApoL6, ApoL6 is a member of ApoL family sharing sequence identity within the amphipathic alpha-helix domain. We show that ApoL6 level in WAT is low in fasting, but is induced upon feeding. Genome-wide association studies (GWAS, https://www.gwascentral.org) identified two single nucleotide polymorphisms (SNPs) of ApoL6 to associate with triglyceride and HDL cholesterol levels. ApoL6 ablation in mice causes a greatly diminished WAT mass with smaller adipocyte size. Conversely, overexpression of ApoL6 increases WAT mass with enlarged adipocytes with higher amount of TAG. We show that ApoL6 robustly inhibits lipolysis. ApoL6 directly interacts with the N-terminal domain of Plin1 to block Plin1-HSL interaction, thereby inhibiting lipolysis in adipocytes.

## RESULTS

### ApoL6 is a LD-associated protein expressed primarily in adipose tissue

RT-qPCR and Northern blotting showed that ApoL6 mRNA levels were highly restricted to adipose tissues, including in WAT such as iWAT and eWAT, and also in BAT although greatly lower (Figure 1A). In adipose tissue, ApoL6 mRNA levels were mainly detected in the adipocyte fraction but not in the stromal vascular fraction (Figure 1B). During 3T3-L1 adipocyte differentiation, as predicted, we could detect decrease in preadipocyte marker Pref-1, and induction of PPARγ and C/EBPα during differentiation. ApoL6 expression was not detected in 3T3-L1 cells prior to adipocyte differentiation, but ApoL6 mRNA and protein levels were increased during adipocyte differentiation (Figure 1C). We also tested human fibroblasts by differentiating them into adipocytes. Similar to that observed in murine 3T3-L1 cells, expression of ApoL6 was increased upon differentiation of human fibroblasts into adipocytes (Figure S1A).

**Figure 1.**
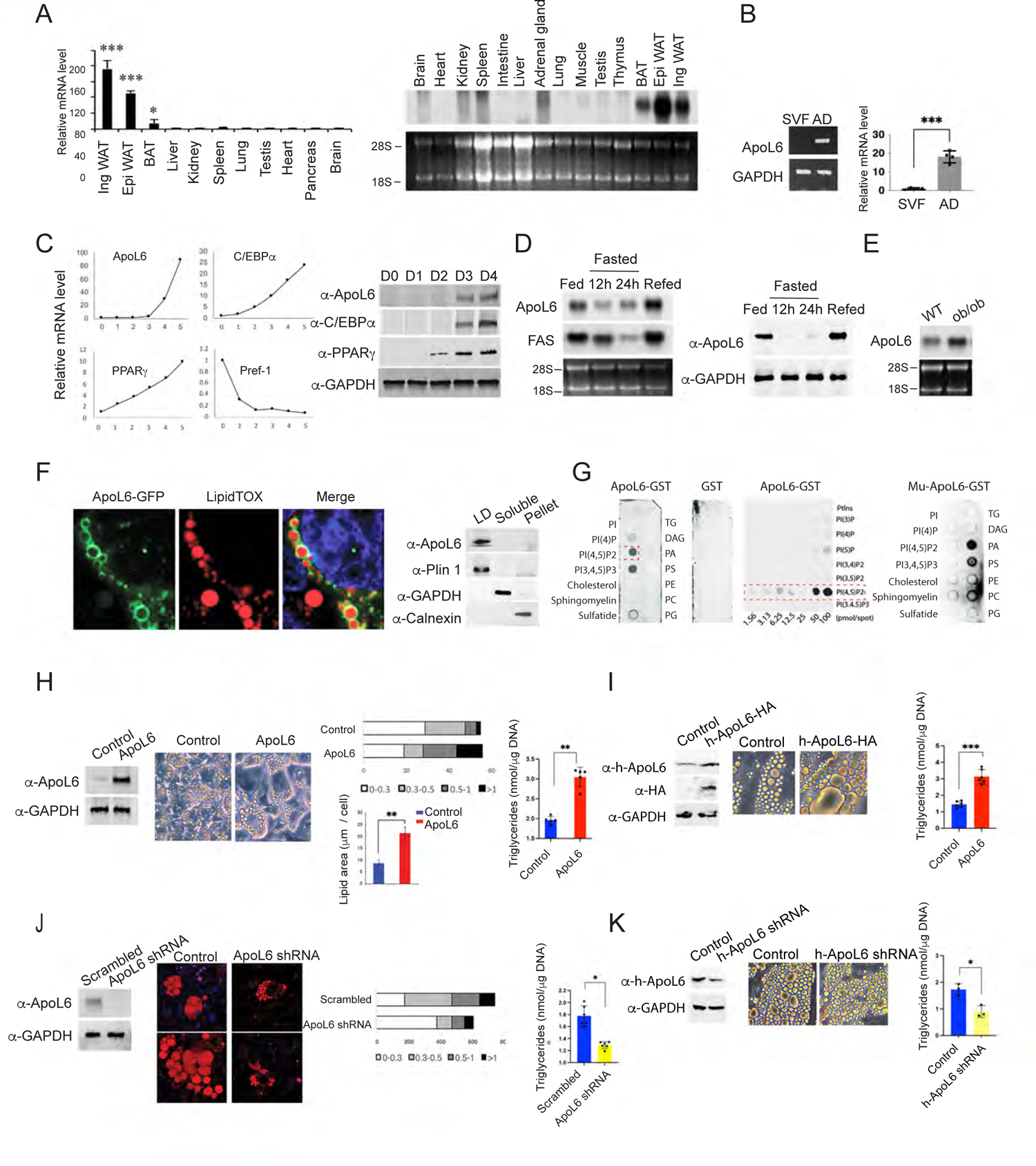
ApoL6 is a LD-associated protein and enhances lipid accumulation. A, RT-qPCR and Northern blotting for ApoL6 mRNA levels in various tissues. B, RT-PCR and RT-qPCR for ApoL6 mRNA levels in stromal vascular and adipocyte fractions. C, RT-qPCR (left) and Immunoblotting (right) during 3T3-L1 adipocyte differentiation. D, Northern blotting (left) and Immunoblotting (right) of WAT tissue. E, Northern blotting of WAT tissue. F, Immunofluorescence after differentiated 3T3-L1cells infected with ApoL6-GFP (green) adenovirus stained with LipidTOX (red) and immunoblotting after sucrose separation (right). G, Immunoblotting after incubation of lipid membrane strips with either WT ApoL6-GST or control GST (right and middle) or mutated ApoL6-GST (Mu-ApoL6-GST, aa286-292, from RYRKLGR to DYDDLGD, right). H, Immunoblotting after infection of adenoviral ApoL6 in differentiated 3T3-L1 adipocytes, quantification of LD size and TAG levels (right). Data represent mean ± SD; \*\**p* <0.01; *n* = 4. Experiment was repeated twice. I, Human adipocytes differentiated from human fibroblasts were infected with lentiviral h-ApoL6-HA, Immunoblotting with h-ApoL6 antibody after infection, Images and TAG levels. \*\*\**p* <0.001; *n* = 4. J, Immunoblotting after scramble and ApoL6-shRNA infection in 3T3-L1 adipocytes. LipidTOX staining, quantification of LD sizes and TAG levels. Data represent mean ± SD; \**p* < 0.05, *n* = 4. Independent experiments were repeat twice. K. Human adipocytes were infected with lentiviral ApoL6 shRNA. Immunoblotting after infection, Images and TAG levels. \**p* <0.05; *n* = 3. Experiments were repeated twice.

We found that ApoL6 mRNA in adipose tissue of mice was decreased upon fasting but increased upon refeeding. In fact, ApoL6 protein in WAT was not detectable in fasting while greatly increased upon refeeding (Figure 1D). ApoL6 mRNA expression was also higher in WAT of ob/ob mice than in non-obese C57BL6 mice (Figure 1E). Overall, the pattern of ApoL6 expression was similar to those genes in lipogenesis that have SRE and E-box motifs where SREBP1c and USF, respectively, are known to bind activate lipogenic genes upon feeding. Indeed, examination of promoter region of ApoL6 gene revealed several E-boxes motifs at −180, −488 and −605, as well as SRE motif at −373. Therefore, we constructed −1 kb ApoL6 promoter-luciferase plasmid and co-transfected with USF-1 or SREBP1c and found ApoL6 promoter activity to be significantly increased upon overexpression of USF-1 and SREBP1c (Figure S1B), indicating their involvement of the ApoL6 induction in the fed condition.

Examination of the amino acid sequence of ApoL6 indicated the presence of an apolipoprotein-like domain in the middle of ApoL6, a hydrophobic domain of potential membrane association near the N-terminus, and the coil-coil domain near the C-terminus. However, ApoL6 lacks N-terminal signal sequence. To examine intracellular localization of ApoL6, a full-length ApoL6-GFP fusion construct was transfected into differentiated 3T3-L1 adipocytes. We detected ApoL6-GFP fluorescence in adipocytes highly localized at LD stained by LipidTOX (Figure 1F left). We also fractionated the adipocyte lysates by centrifugation through sucrose gradient and the fractions were subjected to immunoblotting. As Plin1, ApoL6 was detected mainly in the floating LD fraction, but not in soluble fraction (cytosol) where GAPDH was detected, nor pellet fraction (all membranes and nucleus) where calnexin was detected (Figure 1F right), confirming ApoL6 localization on LD.

Next, to understand how ApoL6 is associated with LD which is composed of core TAG and CE and a surface phospholipid monolayer, we incubated ApoL6-GST (Figure S1C) with membrane strip containing various phospholipid species. As shown in Figure 1G, we detected a selective and strong ApoL6 binding to phosphorylated PIs, such as PI(4)P, PI(4, 5)P2 and PI(3,4,5)P3, but not other phospholipid species. GST control did not show any phospholipid binding. By using varying concentrations of phospholipid species, we found that ApoL6 preferably interacted with PI(4, 5)P2 (Figure 1G middle).

Incubation with various ApoL6 deletion-GST fusion constructs (Figure S1C) showed that C-terminal domain of ApoL6 (aa211-321) was responsible for binding to PI(4,5)P2 (Figure S1B). In fact, we found the consensus amino acid sequence for PIP2 binding, RKLQR (K/H/R)(K/H/R)XX (K/H/R) at aa288-292 of the ApoL6 C-terminal domain. Absence of ApoL6 binding to PI suggested that the negative charges might be important for ApoL6 interaction with PI(4,5)P2. Therefore, we generated ApoL6 constructs containing mutation of amino acids of positive charges, RKLQR, into amino acids of negative charges, CCCLQD (Mu-ApoL6-GST). Indeed Mu-ApoL6-GST did not interact with PI(4,5)P2. However, we detected ApoL6 interaction mainly with PA (Figure 1G right), These observations show the positive charges at the C-terminal domain of ApoL6 are critical for ApoL6 interaction with PI(4, 5)P2. We conclude that PI(4,5)P2 which is a minor yet important component of LD phospholipid monolayer allows ApoL6 recruitment to LD, probably via electrostatic interaction.

### ApoL6 promotes triglyceride and LD accumulation in adipocytes

To understand the ApoL6 function in adipocytes, we overexpressed ApoL6 in 3T3-L1 cells by adenoviral infection (Figure 1H left) and the cells were differentiated into adipocytes. Expression levels of adipogenic transcription factors, such as C/EBPα and PPARγ, as well as lipogenic enzymes, such as FAS and ACC, did not show differences between control and ApoL6 overexpressing cells, indicating neither adipogenesis nor lipogenesis was affected by ApoL6 overexpression (Figure S1D). Remarkably, however, we detected that LD were significantly larger in ApoL6 overexpressing adipocytes compared to control cells (Figure 1H middle). Quantification of lipid area showed a higher lipid staining and content in ApoL6 overexpressing cells (Figure 1H middle). Accordingly, total TAG content in ApoL6 overexpressing cells measured by a biochemical assay was significantly higher than control cells (Figure 1H right). We also employed human adipocytes in documenting ApoL6 effect on TAG and LD accumulation. We infected adipocytes differentiated from human fibroblasts with h-ApoL6 lentivirus (h-ApoL6-HA). ApoL6 protein level was increased by 3-fold upon lentiviral infection (Figure 1I left) and LD size was larger upon h-ApoL6 infection (Figure 1I middle). Accordingly, TAG levels were 2.2-fold higher in h-ApoL6 overexpressing human adipocytes compared to control adipocytes (Figure 1I right).

We next performed ApoL6 knockdown experiments in adipocytes. Infection of lentiviral ApoL6 shRNA greatly decreased ApoL6 expression in 3T3-L1 adipocytes (Figure 1J left). shRNA mediated ApoL6 knockdown caused a significant decrease in LD size (Figure 1J left and middle) and TAG content was decreased by 25% (Figure 1J right). We also performed ApoL6 knockdown experiments in human adipocytes by infecting human shApoL6 lentivirus. ApoL6 expression was decreased by approximately 50% upon shApoL6 lentiviral infection. Similar to that observed in 3T3-L1 adipocytes, the size of LD and TAG levels in human adipocytes were decreased also (Figure 1K). Overall, these results demonstrate that ApoL6 can increase LD size and TAG content in adipocytes.

### ApoL6 inhibits lipolysis in adipocytes

So far, we found that ApoL6 increased TAG and LD content in both rodent and human adipocytes, while not affecting adipocyte differentiation or lipogenic process. Therefore, we hypothesized whether ApoL6 affects TAG hydrolysis, i.e., lipolysis. We examined lipolytic rate by measuring FFA and glycerol release in 3T3-L1 adipocytes after ApoL6 overexpression or knockdown. Indeed, FFA release was significantly lower in ApoL6 overexpressing adipocytes than in control cells in both basal and isoproterenol-stimulated conditions by approximately 30%. Glycerol release was lower in basal and stimulated conditions by 30% and 60%, respectively (Figure 2A). We also tested effect of ApoL6 in human adipocytes. As observed in 3T3-L1 adipocytes, FFA release was significantly lower in h-ApoL6 overexpressing human adipocytes in both basal and stimulated conditions (Figure 2B). These results reveal that overexpression of ApoL6 inhibits lipolysis in adipocytes.

**Figure 2.**
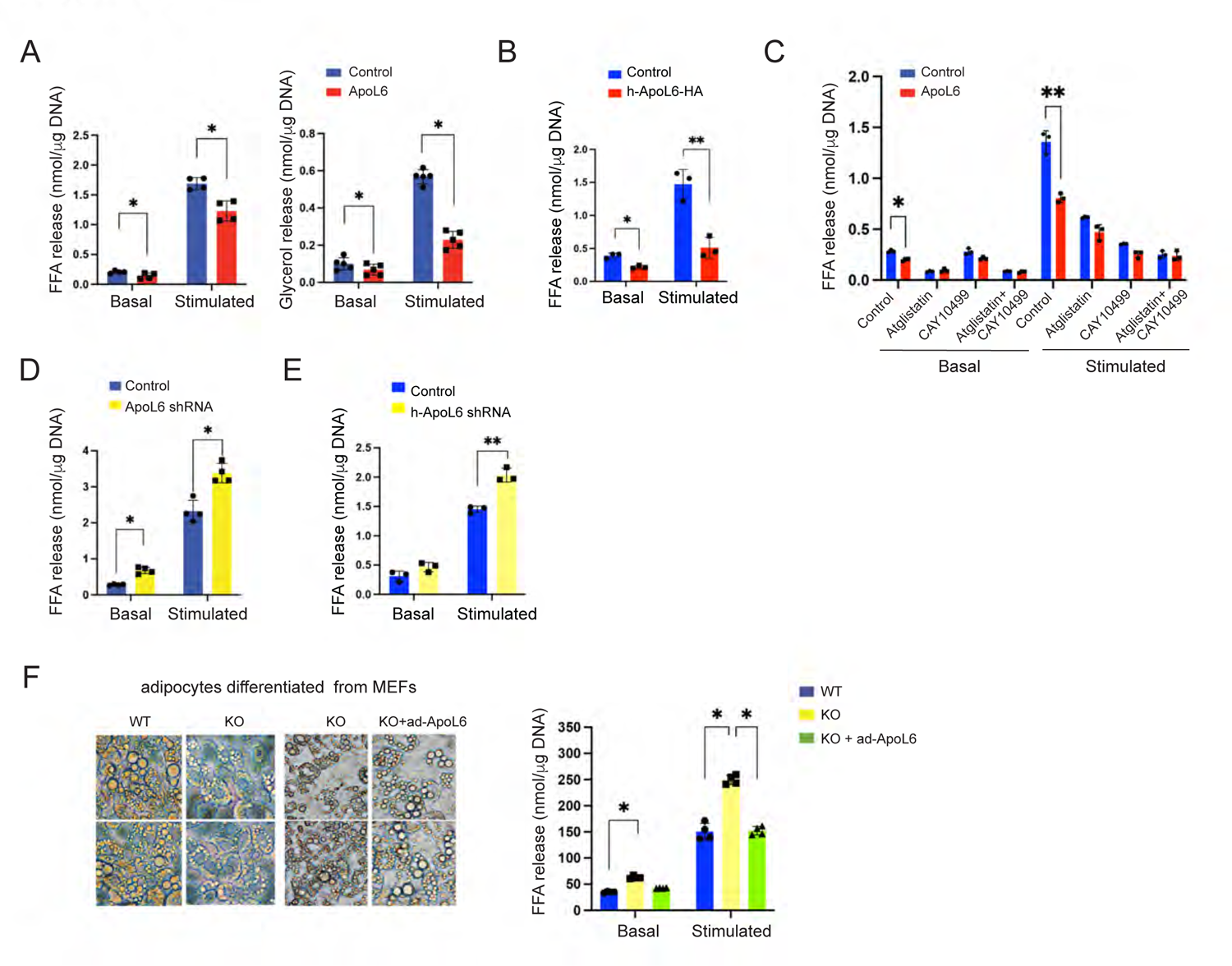
ApoL6 inhibits lipolysis. A, Differentiated 3T3-L1 adipocytes were infected with inducible lentiviral ApoL6, Induced by treating with Dox for 3 hrs before experiment. No Dox-treatment was used as control. FFA and glycerol release in media in basal and isoproterenol treated, stimulated conditions. B, FFA release from human adipocytes after infected with lentiviral h-ApoL6-HA. Data represent mean ± SD; \**p* <0.05; \*\**p* <0.01; *n* = 4. Experiment was repeated twice. C, FFA release from differentiated 3T3-L1 cells cultured with Atglistatin (ATGL inhibitor) and CAY10499 (HSL inhibitor). \**p* <0.05; \*\**p* <0.01; *n* = 3. D, FFA release from differentiated 3T3-L1 adipocytes after ApoL6 shRNA knockdown. E, FFA release from human adipocytes after infected with lentiviral h-ApoL6 shRNA. *p* <0.05; \*\**p* <0.01; *n* = 3. Experiment was repeated twice. F, MEF prepared from WT and ApoL6 KO E13.5 embryos were differentiated into adipocytes and were infected with ApoL6 adenovirus. Image (left) and FFA release (right). Data represent mean ± SD; \**p* < 0.05, *n* = 4. Independent experiments were repeated twice.

When we treated 3T3-L1 adipocytes with ATGL inhibitor, Atglistatin, or HSL inhibitor, CAY10499, as expected, lipolysis was inhibited effectively by these inhibitors. More importantly, after inhibitor treatment, we did not observe any differences in lipolysis between control and ApoL6 overexpressing cells (Figure 2C). These results demonstrate that ApoL6 could not affect lipolysis when ATGL and HSL activities were inhibited. As indicated above, ApoL6 associates with LD by binding to PI(4,5)P2, although ApoL6 with PI(4,5)P2 binding site mutation may still associate with LD via binding to PA. Here, we examined whether ApoL6 binding to PI(4,5)P2 is critical for ApoL6 inhibition of lipolysis. However, overexpression of wild type (WT) ApoL6 and Mu-ApoL6 did not show any differences in the inhibition of lipolysis in both basal and stimulated conditions (Figure S2A). Although future studies on significance of PI(4,5)P2 may be useful. our present results show that ApoL6 inhibition of lipolysis does not require PI(4,5)P2 binding by ApoL6.

Next, we performed ApoL6 knockdown to further examine ApoL6 effect on lipolysis. ApoL6 level was effectively decreased when we transduced ApoL6 shRNA lentivirus in differentiated 3T3-L1 adipocytes. ApoL6 knockdown resulted in a significant increase in lipolytic rate as measured by FFA release in both basal and stimulated conditions (Figure 2D). Similarly, upon ApoL6 knockdown of human adipocytes, FFA release was higher in both basal and stimulated conditions, differences in lipolysis were especially apparent upon isoproterenol treatment, increasing by 40% (Figure 2E). To further verify the loss-of-function effect of ApoL6 on lipolysis, we employed adipocytes differentiated from fibroblasts (MEF) isolated from ApoL6 KO mouse embryos. WT and ApoL6 KO adipocytes did not show any differences in the expression of adipogenic markers, such as C/EBPα, PPARγ, or lipogenic genes, such as FAS and mGPAT (Figure S2B). However, ApoL6 KO adipocytes had noticeably smaller LDs. Moreover, FFA release was significantly higher in ApoL6 KO adipocytes compared to WT adipocytes. Finally, we performed a rescue experiment using these adipocytes differentiated from ApoL6-KO MEF. We infected the differentiated ApoL6 ablated adipocytes with ApoL6 adenovirus that clearly showed expression from ApoL6 adenovirus. Indeed, adenoviral ApoL6 infection in ApoL6-KO MEF derived adipocytes could rescue lipid accumulation and lowered FFA release (Figure 2F right). All together, these results demonstrate that ApoL6 inhibits lipolysis to increase LD size in adipocytes.

### ApoL6 ablation leads to lower WAT mass with higher lipolysis in mice

To evaluate the physiological significance of adipose ApoL6 function in vivo, since ApoL6 was detected primarily in adipocytes of WAT, we generated global ApoL6 knockout mice (ApoL6 KO) by employing the CRISPR-Cas9 system. Guide RNA was designed to target the third exon of ApoL6 gene (Figure 3A upper). By sequencing of the ApoL6 genomic region, we verified one KO mouse line having an 11 bp deletion in the coding region of the third exon (Figure 3A upper), which was germline transmitted (Figure 3A upper). Immunoblotting showed absence of ApoL6 protein in WAT and BAT of ApoL6 KO mice (Figure 3A lower). On normal chow diet, body weight did not show significant differences between WT and ApoL6 KO mice. We therefore subjected ApoL6-KO mice to HFD feeding. ApoL6 KO mice showed significantly lower body weights (BW) after HFD feeding for 7 wks (Figure 3B). WAT depot weights also were significantly reduced in ApoL6 KO mice compared to WT mice, suggesting that ApoL6 KO mice were prevented from HFD-induced obesity (Figure 3B and 3C). H&E staining and imaging, as well as quantification of WAT showed ApoL6 KO mice had smaller adipocyte size compared to WAT of WT mice (Figure 3D). Glucose tolerance test (GTT) and insulin tolerance test (ITT) after 10 wks of HFD feeding indicated that, compared to WT mice which had high glucose levels and insulin resistance as expected, ApoL6 KO mice had significantly lower glucose levels at 15, 30 and 60 min after glucose injection (Figure 3E). These ApoL6 KO mice also exhibited improved insulin sensitivity during ITT (Figure 3E). Serum TAG and total cholesterol levels were significant lower in ApoL6 KO mice than WT mice, although serum FFA levels did not show significant differences (Figure 3F). We then examined lipolysis by using dispersed adipocytes from WAT. Adipocytes from ApoL6 KO mice compared to WT mice showed significant higher FFA and glycerol release (Figure 3G). As expected, in basal condition, treatment with Atglistatin or both Atglistatin and CAY10499 inhibited FFA release in WT and ApoL6 KO adipocytes. In simulated condition, treatment with Atglistatin, CAY10499, or both, significantly decreased FFA release in WT by 14, 72 and 84%, respectively. These inhibitor treatment on ApoL6 KO adipocytes also inhibited lipolysis as in wild type cells (Figure 3H). These results on lipolysis measured in dispersed adipocytes prepared from WAT of ApoL6 KO and WT mice are consistent with the above-described results obtained from cultured adipocytes, demonstrating that absence of ApoL6 increases lipolysis in vivo to prevent diet-induced obesity.

**Figure 3.**
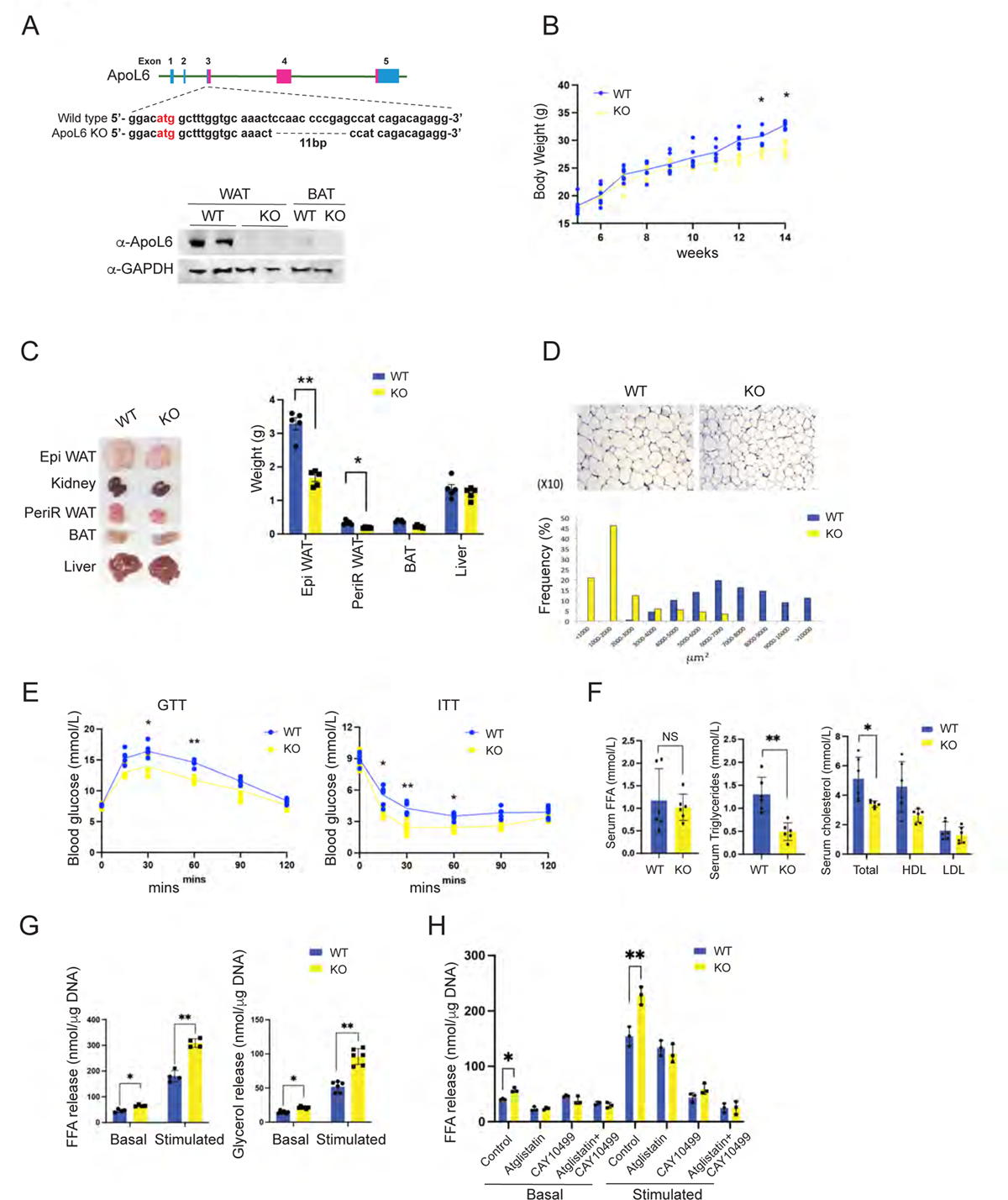
ApoL6 ablation in mice results in lean phenotype with increased WAT lipolysis. A, 11 bp deletion in exon 3 of ApoL6 gene and ApoL6 protein level in WAT and BAT of WT and ApoL6 KO mice. B, BW during HFD feeding. C, Tissue imaging and tissue weights after 10 wks of HFD feeding. D, H&E staining and quantification of cell sizes of WAT. E, GTT and ITT after HFD, n=5, *, p<0.05; **, p<0.01. F, Serum FFA, TAG and cholesterol levels. Data represent mean ± SD; *n* = 6, \**p* < 0.05, \*\**p* <0.01. G, FFA and glycerol release of dispersed adipocytes isolated from WAT. H, FFA release of primary adipocytes cultured with inhibitors. Data represent mean ± SD; \**p* < 0.05, \*\**p* <0.01; *n* = 4. Experiments were repeated twice

### Adipocyte-specific ApoL6 overexpression increases WAT mass and lipid accumulation in mice

In studying ApoL6 function in vivo, next, we generated transgenic mice overexpressing ApoL6 in adipose tissue. We generated transgenic mouse line overexpressing Myc-tagged ApoL6 driven by the −5.4 kb aP2 promoter (aP2-ApoL6 TG) (Figure 4A-4H). ApoL6 transgene expression was examined by RT-qPCR and immunoblotting (Figure 4A and S4A). ApoL6 transgene was expressed specifically in adipose tissue and was barely detectable in other tissues examined in 10-wk old mice (Figure 4A). Immunoblotting using ApoL6 antibody showed that ApoL6 protein levels in aP2-ApoL6 TG WAT was approximately 3-fold higher than that of WT mice.

**Figure 4.**
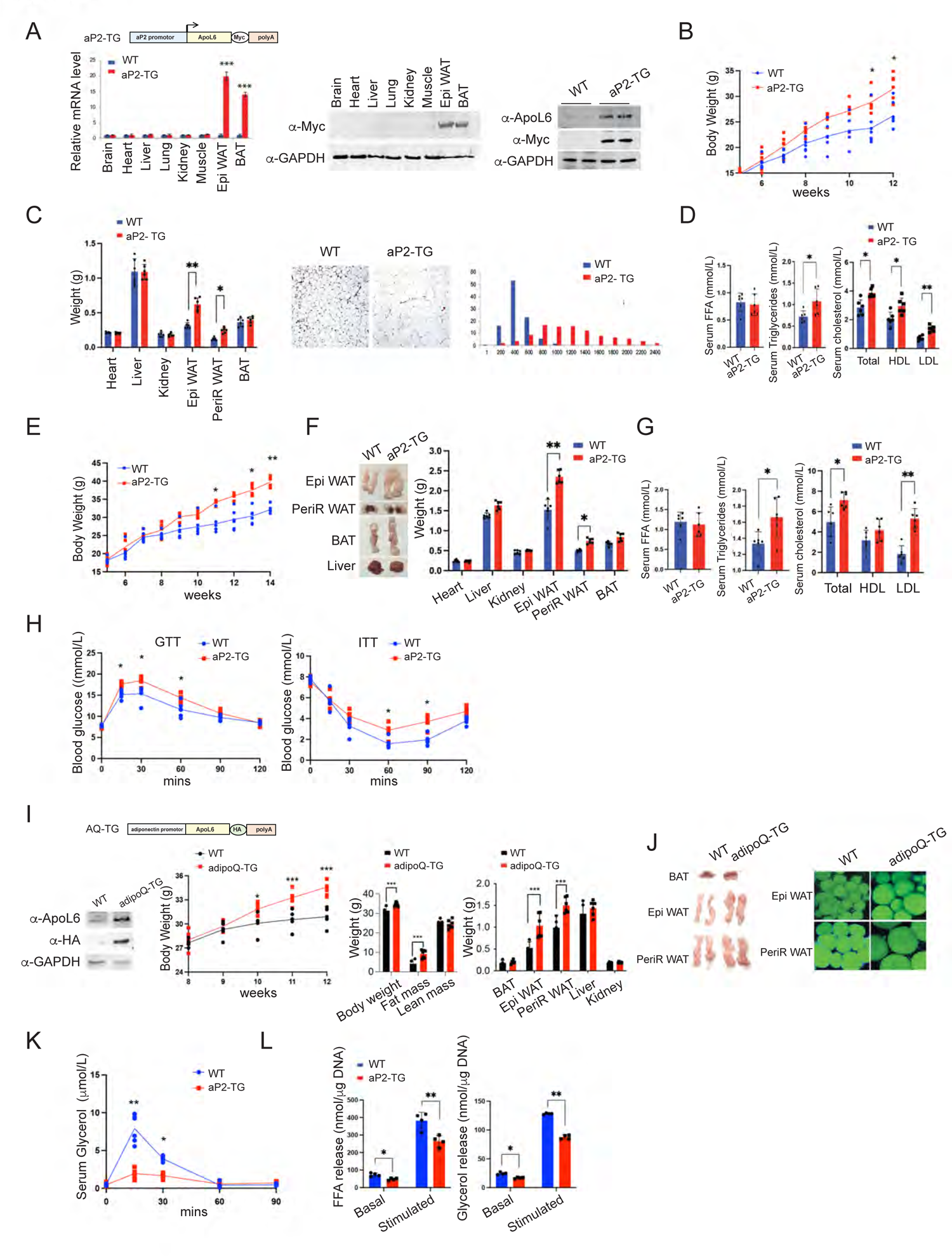
Overexpression of ApoL6 in adipocytes in mice increases WAT mass. A, aP2-ApoL6 TG design (left upper), RT-qPCR (left lower), immunoblotting of transgene expression in different tissues and ApoL6 protein levels in WAT (right). B, BW in chow (right). C, Tissue weights of 24 wk-old mice (left). H&E staining and quantification of cell sizes of WAT (right two panels). D, Serum FFA, TAG and cholesterol levels. E, BW during HFD. F, Tissue image and tissue weight after 12 wks of HFD feeding. Data represent mean ± SD; *n* = 6, \**p* < 0.05, \*\**p* <0.01. G, Serum FFA, TAG and cholesterol levels after HFD feeding, n=6, *, p<0.05, **, p<0.01. H, GTT and ITT after HFD, n=5, *, p<0.05, Experiment was repeated twice. I, adipoQ-ApoL6 TG design. Immunoblotting using lysates of WAT from adipoQ-ApoL6 TG. BW and WAT mass measured by using EchoMRI and tissue weights after 12 wks on HFD. J, Image of tissues and whole mount staining of WAT with LipidTOX green after HFD feeding. K, In vivo lipolysis assay: Serum glycerol levels at different time points after mice were injected with isoproterenol at 10 mg/kg BW. *n* = 6, \**p* < 0.05, \*\**p* <0.01. Experiments were repeated twice. L, FFA and glycerol release from dispersed adipocytes isolated from WAT. Data represent mean ± SD; *n* = 6, \**p* < 0.05, \*\**p* <0.01. Experiments were repeated twice.

When on normal, chow diet, aP2-ApoL6 TG mice at 11-12 wks of age showed significant higher body weights with higher WAT mass, in comparison to their WT littermates (Figure 4B). Liver and other organ weights remained the same. H&E staining of WAT and quantification showed larger adipocyte size in WAT of aP2-ApoL6 TG, compared to WT mice (Figure 4C). No histological differences were detected in BAT of WT and aP2-ApoL6 TG mice (Figure S4B). Gene expression of adipogenic transcription factors, such as C/EBPα and PPARγ, as well as lipogenic genes, such as FAS, ACC and mGPAT, in WAT did not show any differences between WT and aP2-ApoL6 TG mice (Figure S4C), indicating that ApoL6 did not affect adipogenesis or lipogenesis. aP2-ApoL6 TG mice had higher TAG and cholesterol levels, but serum FFA levels did not show differences (Figure 4D). The mice had comparable glucose and insulin tolerance on chow diet when subjected to GTT and ITT (Figure S4D).

We next subjected aP2-ApoL6 TG mice to HFD. Higher BW of aP2-ApoL6 TG mice on HFD compared to WT littermates started to show after 7 wks (Figure 4E). aP2-ApoL6 TG mice had significantly higher WAT mass with larger adipocyte size compared to WT mice (Figure 4F). Serum TAG, cholesterol and LDL/VLDL levels were elevated significantly in aP2-ApoL6 TG mice, although no differences in serum FFA levels were detected (Figure 4G). GTT and ITT showed significantly higher glucose levels at various time points in aP2-ApoL6 TG compared to WT mice (Figure 4H), indicating that aP2-ApoL6 TG mice on HFD were more glucose intolerant and insulin resistant than WT littermates. Since insulin resistance is well associated with hepatosteatosis, we also examined liver lipid accumulation. Interestingly, differ from those mice on chow diet having no histological differences, Oil Red O staining showed higher lipid accumulation and TAG content in livers of aP2-ApoL6 TG compared to WT mice (Figure S4E). Therefore, we also examined ApoL6 expression in liver. ApoL6 was not detected in the livers of 12-wk old chow diet fed mice, but was detected in livers of HFD fed mice, indicating that HFD caused an induction of endogenous ApoL6 expression even in livers of mice when hepatic TAG content was high. We also found that ApoL6 expression was higher in livers of 30 wk-old than 12 wk-old mice (Figure S4F), implicating endogenous ApoL6 might be induced also with aging as hepatic lipid content increased.

Since aP2 promoter is known to be not only active in adipocytes but may also be active in other cell types such as macrophages, we also have generated additional ApoL6 transgenic mouse line overexpressing HA-tagged ApoL6 driven by the −5.2 kb adiponectin promoter (AdipoQ-ApoL6 TG) (Figure 4I and 4J). Similar to aP2-ApoL6 TG mice, ApoL6 protein levels in WAT of adipoQ-ApoL6 TG mice had 3-4 fold higher ApoL6 protein levels than in WT mice. We examined these adipoQ-ApoL6 TG mice after subjecting them to HFD. The adipoQ-ApoL6 TG mice showed significantly higher BW after 10 wks. EchoMRI also showed adipoQ-ApoL6 TG mice having higher BW and WAT mass without changes in lean body mass (Figure 4I). Weights of WAT depots, such as Epi-WAT and PeriR-WAT, but not weights of various other tissues, were significantly higher in adipoQ-ApoL6 TG mice than WT mice after 12 wks of HFD (Figure 4J left). Whole mount staining of WAT with LipidTOX green and quantification indicated that the size of adipocytes of adipoQ-ApoL6 TG mice was larger compared to wildtype mice (Figure 4K right). We conclude that adipoQ-ApoL6 TG mice had similar phenotypes as aP2-ApoL6 TG mice, further confirming that the changes in aP2-ApoL6 TG mice and adipoQ-ApoL6 TG mice are indeed due to overexpression of ApoL6 in adipocytes.

To examine ApoL6 effect on lipolysis in vivo, we next administered nonselective β-agonist isoproterenol to artificially stimulate whole-body lipolysis (Qiao et al. 2011). We then examined blood glycerol concentration at different time points in WT and aP2-ApoL6 TG mice after isoproterenol administration. As expected, WT mice showed a rapid response of lipolysis to isoproterenol treatment, by increasing release of glycerol by 16- and 8-fold than basal condition, 15 and 30 mins after isoproterenol injection, respectively. More importantly, aP2-ApoL6 TG mice had greatly lower increase in glycerol release of only 4-fold higher than basal level at 15 and 30 mins after isoproterenol administration (Figure 4K). These results clearly showed in vivo context a decrease in lipolysis from adipose tissue by ApoL6 overexpression. We also used dispersed adipocytes from WAT of these mice for lipolysis assay in vitro. We detected significantly lower FFA and glycerol release from dispersed adipocytes isolated from WAT of aP2-ApoL6 TG mice than those from WT mice in both basal and stimulated conditions (Figure 4L). Overall, we conclude that ApoL6 inhibits lipolysis in adipose tissue.

### ApoL6 directly interacts with Plin1

To start to understand the mode of action of ApoL6 in inhibiting lipolysis, we investigated ApoL6 interacting protein(s). We identified ApoL6 interacting proteins by immunoprecipitation (IP) with HA antibody using total LD-associated proteins extracted form floating fraction of WAT from adipoQ-ApoL6 TG overexpressing HA-tagged ApoL6, as well as those from ApoL6-HA overexpressing 3T3-L1 adipocytes. The affinity purified proteins using HA antibody beads were subjected to Mass spectrometry (MS) analysis and the experiments were repeated several times. Indeed, proteins involved in lipolysis, such as Plin1, HSL and MGL, were detected by MS analysis multiple times. Since ApoL6 is associated with LD in adipocytes and inhibits lipolysis, we hypothesized that ApoL6 might form a lipolytic complex at the LD surface. We tested our hypothesis by first immunoprecipitating with HA antibody for ApoL6, followed by IP with either HSL or Plin1 antibody. The samples were subjected to native acrylamide gel electrophoresis. Indeed, we detected the same very high molecular weight complex, when the samples of either purified by ApoL6 antibody followed by HSL antibody, or by ApoL6 antibody followed by Plin1 antibody, were loaded (Figure 5A). Potentially, a lipolytic complex containing ApoL6 may be formed on LD in hydrolyzing TAG in adipocytes. Although we did not pursue all components of this complex, MS analysis of this high molecular weight band detected several proteins, including ApoL6, Plin1 and HSL.

**Figure 5.**
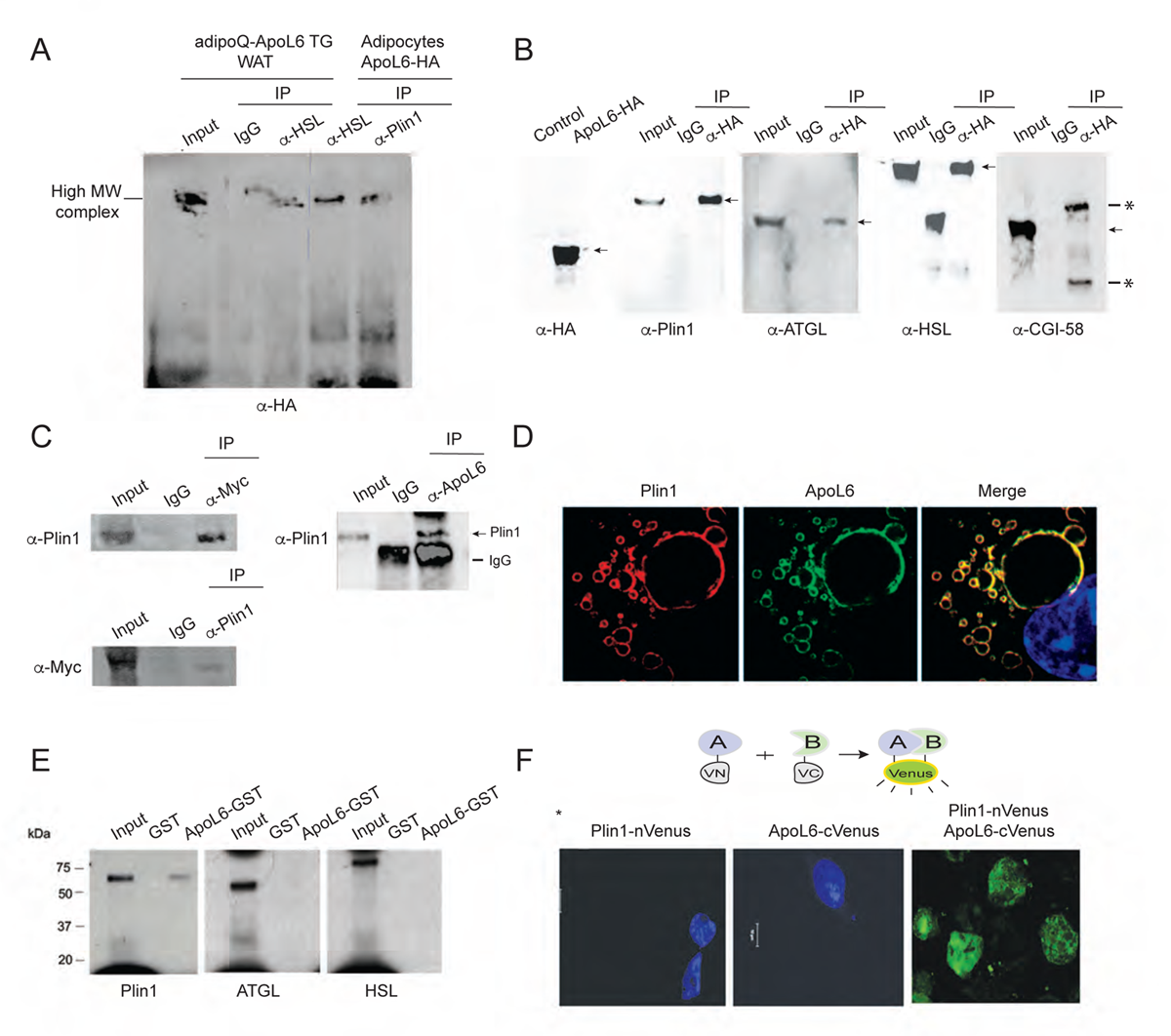
ApoL6 directly interacts with Plin1. A, Total LD-associated proteins from WAT of adipoQ-ApoL6 TG mice and lysates of differentiated 3T3-L1 cells overexpressing ApoL6-HA were first IP with HA antibody, and the elution fractions were subjected to secondary IP with HSL antibody or Plin1 antibody. Immunoblotting with HA antibody after secondary IP using native gel. B, Co-transfection of ApoL6-HA with Plin1, ATGL, HSL and CGI-58 in HEK293 cells. Immunoblotting after lysates were IP with HA antibody. Arrows, positions of interests; *, IgG band. C, Immunoblotting after IP, using lysates of WAT of aP2-ApoL6 TG (left) and WT mice (right). D, Immunofluorescence of differentiated 3T3-L1 adipocytes using Plin1 antibody (mouse) and ApoL6 antibody (rabbit) followed by Alexa fluor plus 594 anti-mouse (read) and Alexa Fluor plus 488 (green) anti-rabbit secondary antibodies, respectively. E, ApoL6-GST was incubated with S35 labeled in vitro transcribed/translated Plin1, ATGL and HSL before GST pull-down. F, Bimolecular Fluorescence Complementation (BIFC) after co-transfection of Plin1 (fused to N-terminus of Venus) and ApoL6 (fused to c-terminus of Venus protein) in HEK293 cells.

We next transfected into HEK293 cells with ApoL6-HA along with Plin1 and HSL, as well as other proteins that are known to be involved in lipolysis, such as ATGL and CGI-58. Co-IP experiments showed ApoL6 interaction with Plin1, HSL and ATGL, but not with CGI-58 (Figure 5B). We could also detect ApoL6 interaction with Plin1 by using WAT. Using total LD-associated protens extracted from aP2-ApoL6 TG WAT overexpressing Myc-tagged ApoL6, we performed IP with Myc antibody followed by immunoblotting with Plin1 antibody. Indeed, we detection ApoL6-Plin1 interaction. Conversely, IP with Plin1 antibody followed by blotting with Myc antibody also detected ApoL6-Plin1 interaction (Figure 5C left). Similar results were obtained when we used WT WAT by IP with ApoL6 antibody for ApoL6 and immunoblotting with Plin1 antibody to detect interaction of endogenous ApoL6 and Plin1 (Figure 5C right). We next performed immunofluorescence experiment for localization. Immunofluorescence using ApoL6 and Plin1 antibodies along with fluorescence conjugated second antibodies showed co-localization of these proteins at the LD surface in differentiated 3T3-L1 adipocytes (Figure 5D), as well as in WAT sections (Figure S5).

While ApoL6 may be present in a large complex composed of lipases and other LD-associated proteins for lipolysis, our next goal was to identify a specific protein that directly interacts with ApoL6. To this end, we performed GST pull-down assay. We incubated ApoL6-GST with [^35^S]-Methionine labeled in vitro translated Plin1, HSL, or ATGL. Indeed, ApoL6 did not show direct interaction with either HSL or ATGL. In contrast, ApoL6 showed a signal when incubated with in vitro translated Plin1, indicating direct interaction of ApoL6 with Plin1, but not with HSL or ATGL (Figure 5E). Next, we employed Bimolecular Fluorescence Complementation (BIFC) assay. We generated expression vector of ApoL6 fused to C-terminus of Venus protein and Plin1 fused to N-terminus of Venus. These plasmids were co-transfected into HEK 293 cells. Indeed, as shown in Figure 5F, we detected ApoL6 interaction with Plin1 by BiFC.

### C-terminal domain of ApoL6 is critical for Plin1 interaction and for inhibition of lipolysis

To examine the domains of ApoL6 and Plin1 that interact, we generated and purified various ApoL6-GST deletion proteins (The same deletion GST constructs that we used for incubation with phospholipid membrane strips in Figure 1)(Figure 6A lower left) and incubated them with purified Plin1. The GST pull-down assay showed that all of the ApoL6-GST deletion proteins showed interaction with Plin1 except those containing deletion of 1/3 of ApoL6 C-terminal domain (constructs #3 and #4) indicating ApoL6 C-terminal domain of aa211-321 is required for interaction with Plin1 (Figure 6A lower right). Next, to identify which domain of Plin1 is responsible for ApoL6 interaction, we generated two Plin1 deletion constructs, N-Plin1 (aa1-280) containing HSL binding domain and C-Plin1 (aa281-517) containing ATGL and CGI 58 binding domain (Figure 6B). The full-length Plin1 (F-Plin1) and its deletion constructs along with full-length ApoL6-HA were overexpressed in HEK293 cells. IP with HA antibody followed by immunoblotting with Plin1 antibody showed that ApoL6 interacted with F-Plin1 or N-Plin1, but not with C-Plin1 (Figure 6B). Overall, we conclude that C-terminal ApoL6 interacts with N-terminal Plin1.

**Figure 6.**
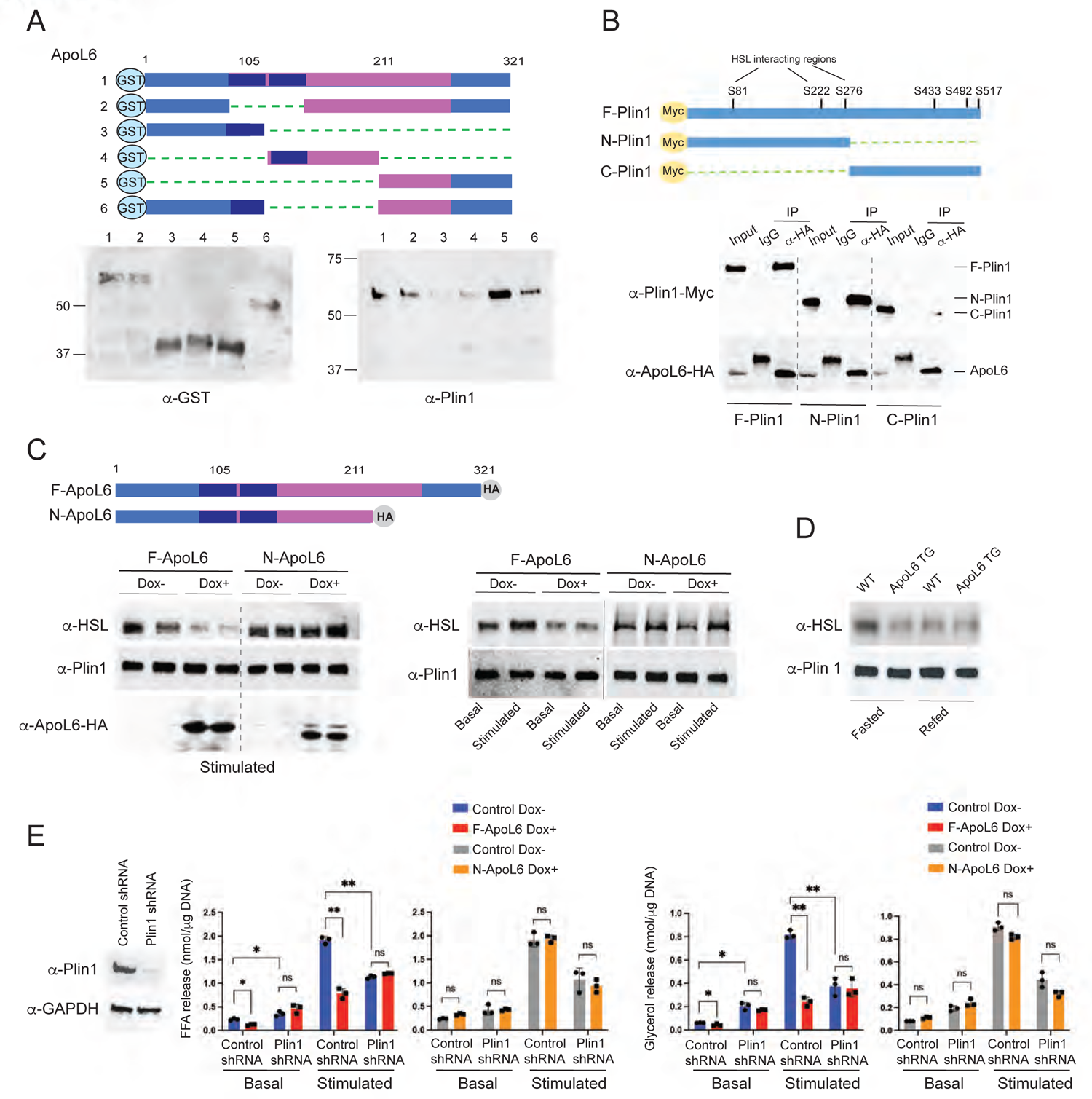
C-terminal domain of ApoL6 is critical for Plin1 interaction to inhibit lipolysis. A, ApoL6-GST deletion constructs (upper), immunoblotting after GST purification (lower left). Purified GST proteins then were incubated with purified full length Plin1. Immunoblotting with Plin1 antibody after GST pull-down (lower right). B, Myc-tagged Plin1 deletion constructs (upper) were co-transfected with F-ApoL6-HA into HEK293 cells. Immunoblotting with Myc antibody after IP with HA antibody. C, Inducible lentiviral F-ApoL6-HA and C-terminal deletion ApoL6 HA (N-ApoL6) (upper) were used to infect HEK293 cells after co-transfection with Plin1 and HSL. Immunoblotting with HSL antibody after IP with Plin1 antibody (left). Immunoblotting with HSL antibody after IP with Plin1 antibody in differentiated 3T3-L1 adipocytes induced to overexpress F-ApoL6-HA and N-ApoL6-HA (right). D, Immunoblotting with HSL antibody after IP with Plin1 antibody using total LD-associated proteins of WAT from wild type and aP2-ApoL6 TG mice in fasted and fed condition. E, Immunoblotting after infecting lentiviral Plin1 shRNA in differentiated 3T3-L1 adipocytes. FFA and glycerol release from 3T3-L1 adipocytes in basal and stimulated conditions. Data represent mean ± SD; \**p* < 0.05, \*\**p* <0.01; *n* = 3. Experiments were repeated twice. F, Schematic model of lipolysis in adipocytes in fed-basal condition and in fasted-stimulated condition.

It is critical for stimulation of lipolysis that Plin1 interacts with HSL that allows HSL translocation from cytosol to LD. To test whether ApoL6-Plin1 interaction has any impact on Plin1-HSL interaction, along with Plin1, we infected lentivirus containing either full-length ApoL6 (F-ApoL6-HA) or ApoL6 deleted of C-terminal domain (aa1-210, N-ApoL6-HA) in a doxycycline (Dox) inducible manner into HEK 293 cells. In the stimulated condition, overexpression of F-ApoL6-HA by Dox treatment compared to control, non-Dox treated cells significantly diminished Plin1-HSL interaction (Figure 6C left), signifying that ApoL6 interaction with Plin1 in turn interrupts Plin1-HSL interaction. In cells infected with ApoL6-N-HA lentivirus, however, no differences in Plin1-HSL interaction were observed whether the cells were overexpressing N-ApoL6 or not (Figure 6C left), demonstrating that ApoL6 C-terminal domain (aa211-321) interaction with Plin1 is required for disruption of Plin1-HSL interaction. We also tested effect of ApoL6 on interaction of endogenous Plin1 and HSL in adipocytes. The 3T3-L1 adipocytes infected with inducible F-ApoL6-HA, were immunoprecipitated with Plin1 followed by immunoblotting with HSL antibody. In non-Dox treated control cells, as predicted, Plin1-HSL interaction was stronger in the stimulated compared to basal condition. In contrast, overexpression of F-ApoL6-HA decreased the Plin1-HSL interaction. Moreover, cells infected with Dox-inducible N-ApoL6-HA, showed no differences whether cells were treated with Dox or not, in either basal or stimulated conditions (Figure 6C right). We next used total LD-associated proteins from WAT to test interaction of endogenous proteins. Upon IP with Plin1, strong Plin1-HSL signal for their interaction was detected from WAT of fasted compared to fed WT mice. However, Plin1-HSL interaction was diminished in WAT of either fasted or fed ApoL6 TG mice (Figure 6D). Taken together, we conclude that, via its C-terminal domain, ApoL6 interacts with Plin1 and the ApoL6-Plin1 interaction prevents Plin1-HSL interaction.

It has been reported that Plin1 phosphorylation at S-81, S-222 and S-276 are required for HSL interaction with Plin1 on LD surface under the stimulated condition (Zhang et al. 2003). To test whether Plin1 phosphorylation affects ApoL6 interaction with Plin1, we generated Plin1 mutation construct by replacing those three Serine residues to Aspartate (Plin1SD) to mimic phosphorylated form. We Incubated ApoL6-GST with either WT Plin1 or Plin1SD purified from 293 FT cells transfected with those expression constructs. Immunoblotting showed that ApoL6-Plin1 interaction was not altered (Figure S6A). We co-transfected HA tagged ApoL6 with either Myc-tagged WT Plin1 or Myc-tagged Plin1SD into 293FT cells and we performed IP with Myc antibody. Immunoblotting with HA antibody showed that ApoL6-Plin1 interaction was not altered when Plin1SD compared to WT Plin1 was used (Figure S6B). Next, we tested whether ApoL6 can inhibit HSL binding to Plin1SD under basal condition. We transfected 293 FT cells with ApoL6 and HSL, along with Myc-tagged Plin1 or Myc-tagged Plin1SD (Figure S6C left). IP with Myc antibody followed by immunoblotting with HSL antibody showed that HSL interaction with WT Plin1 and Plin1SD were not different and ApoL6 prevented Plin1-HSL interaction in either case (Figure S6B right). These results indicate that ApoL6 prevents Plin1-HSL interaction regardless of Plin1 phosphorylation status.

It has been shown that Plin1 is required for stimulated lipolysis (Sztalryd et al. 2003), although Plin1 has been reported to inhibit basal lipolysis (Tansey et al. 2001; Martinez-Botas et al. 2000). For stimulated lipolysis, HSL is phosphorylated and translocated to LD and binds to phosphorylated Plin1 for lipolytic action (Miyoshi et al. 2006). If ApoL6 prevents Plin1-HSL interaction, ApoL6 inhibition of lipolysis should require presence of Plin1. In testing whether Plin1-HSL interaction is required for ApoL6 inhibition of lipolysis, we performed Plin1 knockdown in 3T3-L1 adipocytes by transducing lentiviral Plin1 shRNA (Figure 6E). We then induced expression of ApoL6, before measuring lipolytic rate. In basal condition, consistent with previous reports, Plin1 knockdown significantly increased FFA and glycerol release by 1.6 and 2.3-fold respectively. More importantly, ApoL6 overexpression decreased FFA and glycerol release by 55% and 41%, respectively. This indicates that inhibitory effect of ApoL6 in the basal condition was diminished upon Plin1 knockdown. in the stimulated condition, as expected, FFA and glycerol release were lower by 43% and 54%, respectively, in Plin1 knockdown cells compared to control cells. Moreover, although ApoL6 inhibited lipolysis more than 70% in control cells, ApoL6 could not further inhibit lipolysis in Plin1 knockdown cells. Overall, these data indicate that inhibitory effect of ApoL6 on lipolysis requires presence of Plin1. To better document the role of ApoL6-Plin1 interaction in lipolysis, we next overexpressed N-ApoL6 which does not contain Plin1 interacting domain. Indeed, N-ApoL6 overexpression no longer lowered FFA and glycerol release in both basal and stimulated conditions, regardless of presence of Plin1. These results further confirm that C-terminal domain of ApoL6 is required for its interaction with Plin1, which prevents Plin1-HSL interaction, thereby resulting in inhibition of lipolysis.

In conclusion, as depicted in graphical abstract, in the fed state, ie, non-stimulated basal condition, ApoL6 expression is induced and, the C-terminal domain of ApoL6 interacts with N-terminal domain of Plin1, preventing HSL docking to Plin1. In the fasted state, ie, stimulated condition, decreased ApoL6 levels allow Plin1 to interact with HSL to stimulate lipolysis.

## DISCUSSION

We identify here a new LD-associated protein ApoL6, which is highly restricted to adipose tissue and is induced in fed compared to fasted state to control adipose lipolysis. ApoL6 inhibits lipolysis by directly interacting with Plin1. Thus, ApoL6 knockdown or ablation results in smaller LD with lower TAG content in adipocytes and in WAT of mice that become resistant to diet-induced obesity. Conversely, overexpression of ApoL6 in adipocytes leads to larger LD with high TAG content in adipocytes and in WAT of mice with greater WAT mass, making mice more susceptible to weight gain resulting in impaired glucose tolerance and insulin resistance with ectopic TAG accumulation in the liver.

### ApoL6-Plin1 interaction

We detected various proteins involved in lipolysis, including Plin1, HSL and MGL as a large ApoL6 containing protein complex, a so-called lipolytic complex. Although ApoL6 interacting proteins form a large complex, ApoL6 can directly bind only to Plin1. We also revealed that ApoL6 C-terminal domain is required for the ApoL6-Plin1 binding and that ApoL6 interacts with the N-terminal domain of Plin1. In this regard, ApoL6 contains coiled-coil domain at its C-terminal region, when the coiled-coil domain has been proposed to act as molecular spacer that can either separate functional domains or scaffold large macromolecular complexes (Truebestein and Leonard 2016). In fact, Plin1 is the first and best-studied LD-associated protein (Greenberg et al. 1993; Sztalryd and Brasaemle 2017), In the stimulated state, Plin1 is phosphorylated to release phosphorylated CGI-58. Then, Plin1 N-terminal domain with 3 phosphorylation sites become docking sites for phosphorylated HSL to access LD (Souza et al. 2002). We claim that ApoL6 interacts with the N-terminal domain of Plin1 in order to compete with HSL interaction with Plin1, thereby keeping HSL in a “stand by” status. In this regard, it has been shown that phosphorylation of several specific serine resides at the Plin1 N-terminal domain is required for Plin1-HSL interaction. However, we found that the ApoL6-Plin1 interaction occurs regardless of Plin1 phosphorylation status, preventing HSL-Plin1 interaction. Moreover, we found that ApoL6-C terminal domain which is critical for ApoL6-Plin1 interaction binds to PI(4,5)P2 to be associated with LD. However, mutation of ApoL6 for ApoL6-PIP2 still could associate with LD by an alternate component of phospholipid monolayer and can inhibit lipolysis, suggesting that ApoL6-Plin1 interaction itself might play the major role in ApoL6 inhibition of lipolysis.

### ApoL6 suppression of adipose lipolysis

Our in vitro and in vivo studies reveal the importance of ApoL6 function to inhibit lipolysis. Plin1 is known to have dual function by acting to augment stimulated lipolysis, while suppressing basal lipolysis: Thus, several studies have shown that Plin1 knockdown decreases lipolysis at stimulated condition, indicating the requirement of Plin1 for stimulated lipolysis (Sztalryd et al. 2003). Expression of ApoL6 in the fasted state is suppressed via transcriptional mechanism, and absence of ApoL6 allows interaction of phosphorylated HSL to Plin1 which is phosphorylated at the N-terminal region. Artificially overexpression of ApoL6 competes with HSL binding to Plin1 to inhibit lipolysis. Deletion of ApoL6 C-terminal domain or KD of Plin1 diminished ApoL6 inhibitory effect on lipolysis demonstrating the importance of ApoL6-Plin1 binding for ApoL6 function. Under the basal or fed condition, however, Plin1 plays a crucial role in restricting adipose lipolysis. In *vivo*, and *in vitro* studies, including those on Plin1-null mice, have shown that reduced levels of Plin1 or absence of Plin1 results in increased basal lipolysis (Miyoshi et al. 2010; Martinez-Botas et al. 2000; Tansey et al. 2001). Pin1 binds CGI58 with high affinity and thereby suppresses CGI-58-ATGL interaction (Granneman et al. 2009). C-terminal domain of Plin1 has been shown to be essential for binding to CGI-58, and this interaction stabilizes CGI-58 localization on the LD (Gandotra, Lim, et al. 2011). Adipocytes overexpressing mutant Plin1 exhibited elevated basal lipolysis. Thus, expression of Plin1 frameshift mutations that lack C-terminal amino acid sequence or Plin1 with C-terminal truncation, could reduce TAG storage in cells by increasing TAG turnover (Gandotra, Le Dour, et al. 2011). Here, we show that, in the fed condition, ApoL6 that is transcriptionally activated involving USF and SREBP1c, increased ApoL6 levels to inhibit lipolysis by blocking interaction of N-terminal domain of Plin1 to HSL In the basal condition, artificial overexpression of ApoL6 did not increase basal lipolysis, since ApoL6 interacting domain of Plin1 resides at N-terminus, but not C-terminus where CGI-58 interacts. Future studies are needed to understand how ApoL6 could inhibit lipolysis in basal condition in cultured adipocytes. Physiologically, although ApoL6 decreases during fasting, overexpression of ApoL6 in transgenic mice had greater inhibitory effect on lipolysis in stimulated condition. We propose that ApoL6 protein levels decrease in fasted condition, potentially involving ApoL6 protein degradation mechanism, to allow Plin1-HSL interaction, resulting in stimulation of lipolysis.

### Effect of ApoL6 on adiposity, insulin resistance and FLD

Our global ApoL6 mice showed protection from diet-induced obesity, and mice overexpressing ApoL6 in adipose tissue showed exacerbated adiposity. Accordingly, ApoL6 knockout mice showed improved glucose tolerance and insulin sensitivity, while ApoL6 overexpressing mice had impaired glucose tolerance and insulin resistance. These changes in insulin sensitivity reflect the inhibitory effect of ApoL6 on TAG accumulation in adipose tissue. In this regard, Plin1 KO mice (Tansey et al. 2001; Sohn et al. 2018; Martinez-Botas et al. 2000) had less adipose mass and resistant to diet-induced obesity, but develop glucose intolerance and insulin resistance showing higher serum TAG but normal serum FFA levels (Sohn et al. 2018). Although the authors did not examine liver TAG accumulation, fatty liver disease (FLD) might have contributed to insulin resistance in these mice. Human Plin1 deficiency in heterozygous frameshift mutations had partial lipodystrophy with dyslipidemia and insulin resistant diabetes (Gandotra, Le Dour, et al. 2011). Interestingly, however, Plin1 overexpression in adipose tissue has also been reported to protect mice against diet-induced obesity with inhibited basal and stimulated lipolysis and improved glucose tolerance (Miyoshi et al. 2010). These conflicting results between Plin1 KO and Plin1 overexpressing mice might due to transgene expression in brown adipose tissue and metabolism (Miyoshi et al. 2010).

In humans, Maximum enzymatic activity of HSL was reported to decrease by approximately 40% in 10 patients with familial combined hyperlipidemia and insulin resistance (Reynisdottir et al. 1997). However, controversial results were reported with HSL-deficient mice: some HSL-deficient mouse models were shown to develop FLD (Roduit et al. 2001; Mulder et al. 2003), Adipose-specific deletion of HSL caused FLD by 8 months of age (Xia et al. 2017). Other mouse models were reported to have a low hepatic TAG content (Haemmerle et al. 2002; Voshol et al. 2003; Donnelly et al. 2005). However, loss of function of HSL by mutations in humans was to bring FLD (Fischer et al. 2007; Haemmerle et al. 2006; Albert et al. 2014). Interestingly, we detected ApoL6 in livers of mice on HFD or upon aging. Although ApoL6 is present mainly in WAT but not in liver of young mice, It is possible that increased TAG accumulation in liver upon HFD feeding may have induced ApoL6 expression, contributing to glucose intolerance and insulin resistance. The Human Protein Atlas (https://www.proteinatlas.org) indicated presence of ApoL6, not only in WAT, but also in liver, while we found ApoL6 primarily in adipose tissue of young mice on normal chow diet. This discrepancy might be due to diet and aging. Considering GWAS data of ApoL6 SNP association with triglyceride levels in humans, our studies suggest that ApoL6 may provide a future therapeutic target against obesity/diabetes and FLD.

## METHODS

### Resources

#### Mouse models

ApoL6 transgenic mice: Two strains of ApoL6 overexpressing mice were generated, one was Myc-tagged ApoL6 driven by −5.4 kb aP2 (FABP4) promoter (aP2-ApoL6 TG), the other was HA-tagged ApoL6 driven by the −5.2 kb adiponectin promoter (adipoQ-ApoL6 TG). Mice were genotyped using tail from the mice. Positive mice were mated with C57BL/6J mice and both lines were germline transmitted. All transgenic mice and littermates were on C57BL/6J background. aP2-ApoL6 TG genotype primers were forward 5’-AAACCCAACAGAGCTTGCAG-3’ and reverse 5’-CAAGTCCTCTTCAGAAATGAG-3’. adipoQ-ApoL6 TG genotype primers were forward 5’-TGTCAATTTCAGGGCTCAGGATA-3’ and 5’-GCTTTCGTCAATGTGGTCTGC-3’.

Global ApoL6 knockout mice (ApoL6 KO) were generated by using CRISPR-Cas9 system, targeting translation start site gRNA (TCTGTCTGATGGCTCGGGGTTGG) and Cas9 mRNA were injected into the zygotes. By sequencing the ApoL6 genomic region of KO cells, we verified one KO line having a 11bp deletion in the coding region of the third exon (18 bp after translation start site) and the line was germline transmitted (Figure 2A upper). Generation of mice was performed by the UC Berkeley Gene Targeting Facility. Primers for wild type were forward 5’-AAACTCCAACCCCGAGCCAT-3’ and reverse 5’-GCCCCAATTAAACGCTTTCC-3’; ApoL6 KO mutant primers were forward 5’-GGTGCAAACTCCATCAGA-3’ and reverse 5’-GCCCCAATTAAACGCTTTCC-3’.

We provided either a standard chow diet (Harlan Teklad LM-485) or a high-fat diet (HFD; 45% of calories from fat, 35% of calories from carbohydrates, and 20% of calories from protein; Research Diets) *ad libitum* after weaning. For fasting/feeding experiments, mice were fasted overnight and then fed a high carbohydrate, fat-free diet for 8-16 hrs. High carbohydrate diet (70 kcal% carbohydrates) and High fat diet (45 kcal% fat) were from Research Diets. The light was turned on at 7 a.m. and off at 7 p.m. Animal housing and all protocols and the experimental procedures were approved by the University of California at Berkeley Animal Care and Use Committee.

#### Cell culture and adipocyte differentiation

All cells were cultured at 37°C in high-glucose Dulbecco’s modified Eagle’s medium (Gibco BRL) supplemented with 10% (v/v) heat-inactivated fetal bovine serum (Omega) and 100 U/ml penicillin/streptomycin (Gibco-BRL).

MEF were prepared from E13.5 embryos as described previously (Wang and Sul, 2006). Briefly, Embryos were removed and separated from maternal tissues and yolk sacs were finely minced, digested with 0.25% for 30 min at 37°C, and centrifuged for 5 min at 1,000 × *g*. The pellet was resuspended in culture medium before plating. All differentiation experiments were carried out using cells at passage 2. For adipocyte differentiation of MEF, MEF were split into six-well plates and cultured to confluence. Two days later, the medium was replaced with differentiation induction medium containing 0.5 mM methyl-isobutyl-xanthine (MIX), 1 μM dexamethasone (DEX), 10 μg/ml insulin, 10 μM troglitazone, and 10% (v/v) FBS. MEF were treated with differentiation agents for 4 days, and the medium was renewed every other day.

For 3T3-L1 adipocyte differentiation, 3T3-L1 cells were split into six-well plates and cultured to confluence. Two days later, the medium was replaced with differentiation induction medium containing 0.5 mM MIX, 1 μM DEX, 10 μg/ml insulin and 10% (v/v) FBS for 2 days and changed back to complete media for additional two days.

#### Adenovirus and lentivirus generation and transduction and plasmid transfection

Adenovirus were generated using ApoL6 full length plasmid by Vector Biolabs. Lentiviral construct was made by subcloning ApoL6 and its mutant into pLenti6/V5-D-TOPO vector and inducible expressing vector, pInducer 20 (Addgene). ApoL6 was also subcloned into pcDNA3.1. Lentiviral Plin1 shRNA plasmid was purchased from Sigma. Lentivirus was first packaged in 293 FT cells using co-transfection with LV-MAX Lentiviral packaging mix (ThermoFisher) for 48 hrs, and cells were infected with harvested lentivirus for 48 hrs before further experiments. The 293FT cells were transfected with-444-FAS-Luc along with various expression plasmids, including USF-1 and SREBP1c, by using Lipofectamine 2000 (Invitrogen). Luciferase assays were performed using Dual-Luc reagent (Promega).

#### Bimolecular Fluorescence Complementation (BiFC) assay

BiFC is based on fluorescence detection upon complementation between nonfluorescent fragments, when brought together by an interaction between the two proteins each fused to N- or C-fragment, become fluorescent. We used Venus system (Addgene) by fusing VN210 (Venus aa1-210) with Plin1 (Perilipin 1-nVenus) and VC210 (Venus aa210-238) with ApoL6 (ApoL6-cVenus) and the two constructs were transfected into differentiating 3T3-L1 adipocytes, and the interactions were visualized by fluorescence imaging.

#### Northern blot analysis, RT-PCR and RT-qPCR

Total RNA prepared with Trizol (Invitrogen) were subjected to agarose-formaldehyde gel electrophoresis and blotted onto Hybond N membranes (Amersham). The membranes were hybridized with ^32^P-labeled cDNAs for mouse ApoL6 or FAS. Primer sets used for RT-PCR to produce the cDNA probes were described in supplemental data section. Hybridization was performed as described previously (Wang and Sul, 2006). For RT-qPCR, 1 μg of total RNA were reverse transcribed using SuperScript II (Invitrogen). The cDNAs were mixed with SYBRTM green PCR master mix (Invitrogen), specific primers for ApoL6. Pref-1, C/EBPα and PPARγ. Primer sets: ApoL6 (Pref-1 (5′-GACCCACCCTGTGACCCC-3′ and 5′-CAGGCAGCTCGTGCACCCC-3′), C/EBPα (5′-TGGACAAGAACAGCAACGAG-3′ and 5′-AATCTCCTAGTCCTGGCTTG-3′), PPARγ2 (5′-ACTCTGGGAGATTCTCCTGTTGAC-3′, and 5′-ACTCTGGGAGATTCTCCTGTTGAC-3′, ′), FAS (5′-TGCTCCCAGCTGCAGGC-3′ and 5′-GCCCGGTAGCTCTGGGTGTA-3′) and GAPDH (5′-CATCACCATCTTCCAGGAGCG-3′ and 5′-TGACCTTGCCCACAGCCTTG-3′). Data were analyzed by using ABI7900. Gene expression levels were calculated by normalization to glyceraldehyde 3-phosphate dehydrogenase (GAPDH) by the ΔΔ*C_T_* method. The mean cycle threshold (*CT*) was converted to a relative expression value by the 2^−ΔΔ*CT*^ method, and the range was calculated by the 2^−(ΔΔ*CT* + standard deviation of ΔΔ*CT*)^ method.

#### Immunoblotting

Total lysates were subjected to 8% or 10% SDS-PAGE, transferred to Protran membranes. After blocking with 4% nonfat dry milk in Tris-buffered saline-Tween buffer, the membranes were probed with first antibodies and followed by a horseradish peroxidase (HRP)-conjugated secondary antibody (Bio-Rad). Blots were visualized by enhanced chemiluminescence (PerkinElmer), and images were captured with Bio-Rad ChemiDoc imaging system.

#### Isolation of LD-associated proteins

LD-associated proteins were isolated by using single centrifugation method (Harris et al. 2012). Briefly, 50-100 mg epididymal fat from mice with 800 ul lysis buffer (10mM HEPES and 1mM EDTA, PH 7.4) was minced with a razor blade and homogenized. 200 ug of 60% (w/w) sucrose was added to the tissue sample and incubated on ice for 20 min and mix well. 600 ul lysis buffer with food dye (2ul/1 ml of lysis buffer) was carefully layered on top of the homogenate and centrifuged for 2 hrs at 20,000 g at 4 °C. the tube was frozen thoroughly, and associated proteins were harvested from top layer with RIPA buffer containing phosphatase inhibitor cocktail (Cell Signaling), and protease inhibitor cocktail (Sigma).

#### Immunoprecipitation and GST pull-down

Total LD-associated proteins from differentiated 3T3-L1 cells in buffer containing 1% Triton X-100, 150 mM NaCl, 10% glycerol, 25 mM Tris [pH 7.4], phosphatase inhibitor cocktail (Cell Signaling), and protease inhibitor cocktail (Sigma) were incubated with antibodies against Plin1 at 4°C overnight, followed by the addition of 40 μl of protein A/G-agarose beads (Santa Cruz) and incubation for 1 h. Beads were collected by pulse centrifugation and washed five times with lysis buffer and heated to 99°C for 5 min in 5× SDS sample buffer. The input and the immunoprecipitated fraction were subjected to SDS-PAGE. HSL, Plin1, and GAPDH were detected by immunoblotting with each antibody.

For GST pull-down, bacterially expressed GST-ApoL6 fusion protein on glutathione-agarose beads (Santa Cruz) were incubated with ^35^S labeled proteins made by using TNT Quick Coupled transcription/Translation System (Promega) with ^35^S labeled methionine. The proteins were separated by SDS-PAGE before autoradiography.

#### Lipolysis assay in cultured adipocytes and in mice

12 well plates of differentiated 3T3-L1 adipocytes were washed with Krebs Ringer buffer (KRB; 12 mM HEPES, 121 mM NaCl, 4.9 mM KCl, 1.2 mM MgSO_4_, 0.33 mM CaCl_2_) and incubated at 37°C in 300 μl of Krebs Ringer buffer containing 2% FA-free bovine serum albumin (BSA) and 0.1% glucose in the presence or absence of 10 μM isoproterenol (Sigma) for 1hr.

Primary adipocytes were isolated from epi-WAT after digested at 37°C for 1 hr with collagenase (Roche) in KRB supplemented with 3 mM glucose and 1% FA-free BSA. Digestion products were filtered through nylon mesh and centrifuged. Adipocytes were collected from the upper phase. Cells were then incubated at 37°C in 500 μl of KRB in the presence or absence of 10 μM isoproterenol (Sigma) for 1hr or with 50 μM ATGL inhibitor, Atglistatin (Sigma), and 10 μM HSL inhibitor CAY10499 (Cayman). Lipolysis was assayed by the release of FFAs and glycerol into the media as described previously using the NEFA-HR (Wako, #997-76491) and glycerol reagent (Sigma), respectively.

For In vivo lipolysis, mice were injected intraperitoneally with isoproterenol in PBS at 10 mg/kg. Blood samples from tails were collected at 0, 15, 30, 60 and 90 mins after injection. Serum was subjected to glycerol measurement.

#### Serum metabolite measurements

Blood was centrifuged at 4°C for 15 min for fractionation. TAG levels were measured with the Infinity triglyceride reagent (Thermo-Fisher), and non-esterified FAs (NEFAs) were measured with the NEFA-HR kit (Wako). TAG contents were measure with Triglyceride quantification assay kit (Abcam, #ab65336). Total cholesterol and HDL and LDL cholesterol were measured with Cholesterol assay kit (Abcam ab65390).

#### Protein-lipid Overlay Assay

Membrane lipid strips ((p-6002) Echelon biosciences) were blocked in buffer containing PBS, 0.1% Tween 20, 3% BSA for 1 hr. The strips were then probed with ApoL6-GST for 4 hrs. The strips were washed 3 times with buffer containing PBS and 0.1% Tween 20 and then probed with GST antibody followed by anti-rabbit HRP conjugate as the secondary antibody. The strips were developed with Pierce ECL detection kit (Thermo-Fisher).

#### Adipocyte size determination

Epi-WAT samples were fixed in 10% buffered formalin, embedded in paraffin, cut into 8-μm-thick sections, and stained with hematoxylin and eosin. Adipocyte size was determined with NIH ImageJ software by measuring a minimum of 300 cells per slide, five slides per sample, and three or four mice per group.

#### Statistical analysis

The results are expressed as means ± the standard errors of the means (SD). Student’s *t* test was used for comparisons of two groups. Unless specifically stated otherwise, all significance levels were set at *P* < 0.05. All experiments were repeated at least twice, and representative data are shown.

## Author contributions

Wang Y: designed and performed most of the experiments, analyzed the data and wrote the manuscript.

Nguyen HP: performed adipoQ-ApoL6 TG phenotyping and imaging and lipolysis assays in 3T3-L1 adipocytes.

Xue P: ApoL6 interaction with Plin1 and Plin1 knockdown experiments.

Xie Y: 3T3-L1 cell culture and differentiation into adipocytes.

Yi D: Co-localization of ApoL6 and Plin1.

Lin F: lipolysis assays and mice maintenance. Viscarra JA: generation adipoQ-ApoL6 TG mice

Nnejiuwa U. lbe: Discussion and preparation of the manuscript

Duncan RE: generation of ApoL6 expression constructs.

Sul HS: conceived and devised experiments, supervised the study and wrote the manuscript.

All authors commented on and edited the manuscript.

## SUPPLEMENTAL FIGURES

**Figure S1**

A, Location of E box and SRE motifs at ApoL6 promoter region (upper) and relative luciferase activity after ApoL6 promoter-luciferase construct upon co-transfection with USF-1 and SREBP-1c. *, p<0.05 compared to cells transfected with empty control vector. *, p<0.05, n=4. Experiments were repeated three times. B, Coomassie blue staining of purified GST-tagged full-length ApoL6 and its deletion constructs (upper). Immunoblotting with GST antibody after incubation of lipid membrane strips with different deletions of ApoL6-GST. C, RT-qPCR for expression of adipogenic transcription factors and lipogenic enzymes in differentiated 3T3-L1 adipocytes after infection with control or ApoL6 adenovirus. n=4, Experiments were repeated twice. D, Human fibroblasts were differentiated into adipocytes. RT-qPCR for human ApoL6 mRNA levels before and after differentiation at Day 4. ***, p<0.001, n=3.

**Figure S2**

A, FFA and glycerol release in media in basal and isoproterenol treated conditions after differentiated 3T3-L1 adipocytes were infected either with lentiviral ApoL6 or lentiviral Mu-ApoL6. Data represent mean ± SD, n=6. Experiments were repeated twice. B, RT-qPCR for expression of adipogenic transcription factors and lipogenic enzymes after differentiation of MEF into adipocytes. Data represent mean ± SD; n=6, experiments were repeated twice.

**Figure S4**

A, Immunoblotting with Myc antibody for ApoL6-Myc protein levels in WAT and BAT from aP2-ApoL6 TG mice. B, H&E staining of livers from 20 wk-old mice on chow diet. C, RT-qPCR of adipogenic transcription factors and lipogenic enzymes in WAT. n=4. D, GTT and ITT for mice on chow diet. n=6, experiments were repeated twice. E, Oil Red O staining and TAG levels in livers of mice on HFD. F, RT-PCR for ApoL6 expression in livers of mice.

**Figure S5**

A, Fluorescence imaging of whole mount WAT and immunoblotting with Plin1 (mouse) and ApoL6 (rabbit) antibodies followed by Alexa fluor 594 anti-mouse and Alexa fluor 488 anti-rabbit antibodies, respectively.

**Figure S6**

A, S81, S222, S276 of Plin1 were all mutated to Aspartic acid (Plin1SD). Purified WT Plin1 and Plin1SD were incubated with purified ApoL6-GST. Immunoblotting with GST antibody after IP with Myc antibody (left). B, Co-transfection of ApoL6-HA with Myc tagged WT Plin1 and Plin1SD into 293FT cells. Immunoblotting with HA antibody after IP with Myc antibody. C, Co-transfection of ApoL6-HA, HSL, with Myc tagged either WT Plin1 or Plin1SD into HEK293FT cells (left). Immunoblotting with HSL antibody after IP with Myc antibody (right).

