## Supplementary figures and images for "A new LD protein, ApoL6 disrupts the Perilipin 1-HSL interaction to inhibit lipolysis"

### Supplemental figures

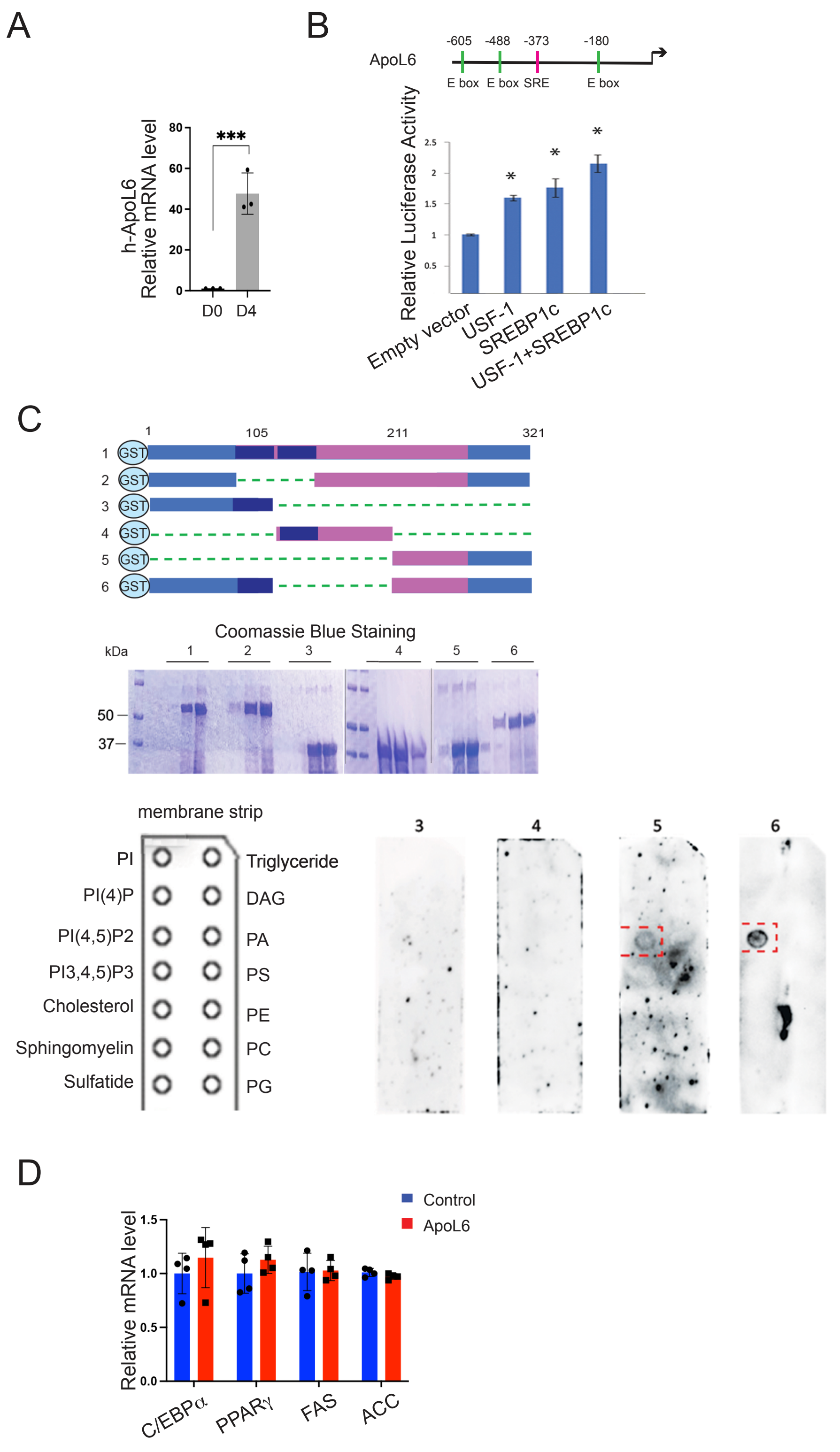

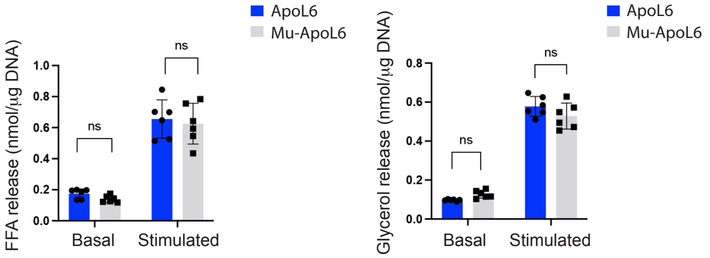

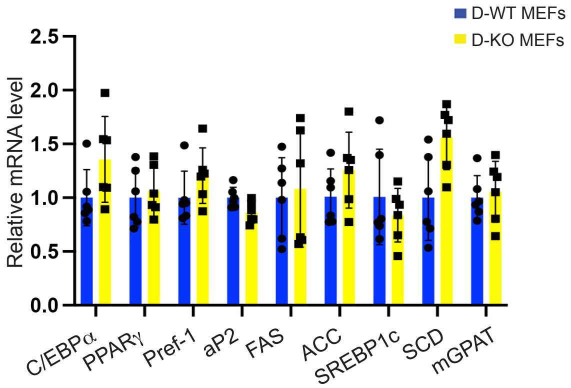

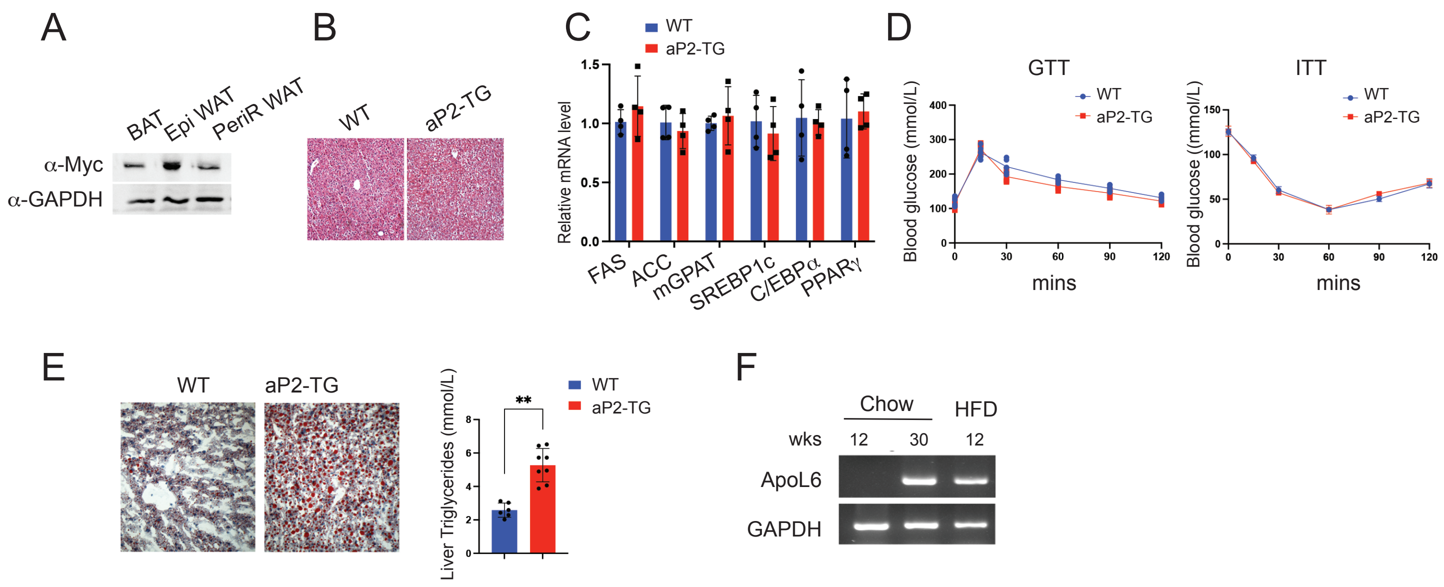

#### Figure S5

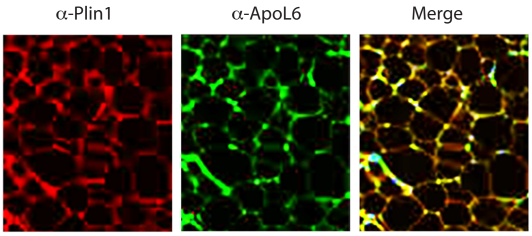

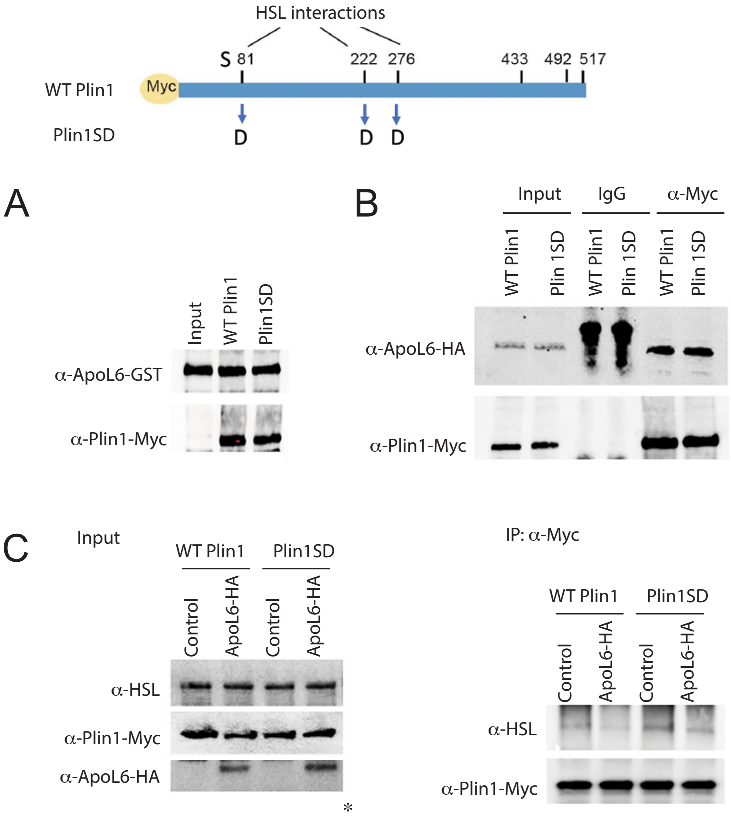
